# Accurate microRNA annotation of animal genomes using trained covariance models of curated microRNA complements in MirMachine

**DOI:** 10.1101/2022.11.23.517654

**Authors:** Sinan Uğur Umu, Vanessa M. Paynter, Håvard Trondsen, Tilo Buschmann, Trine B. Rounge, Kevin J. Peterson, Bastian Fromm

## Abstract

The annotation of microRNAs, an important class of post-transcriptional regulators, depends on the availability of transcriptomics data and expert knowledge. This led to a large gap between novel genomes made available and high-quality microRNA complements. Using >16,000 microRNAs from the manually curated microRNA gene database MirGeneDB, we generated trained covariance models for all conserved microRNA families. These models are available in MirMachine, our new tool for the annotation of conserved microRNA complements from genomes only. We successfully applied MirMachine to a wide range of animal species, including those with very large genomes, additional genome duplications and extinct species, where smallRNA sequencing will be hard to achieve. We further describe a microRNA score of expected microRNAs that can be used to assess the completeness of genome assemblies. MirMachine closes a long-persisting gap in the microRNA field facilitating automated genome annotation pipelines and deeper studies on the evolution of genome regulation, even in extinct organisms.

**Highlights:**

- An annotation pipeline using trained covariance models of microRNA families
- Enables massive parallel annotation of microRNA complements of genomes
- MirMachine creates meaningful annotations for very large and extinct genomes
- microRNA score to assess genome assembly completeness

Graphical abstract

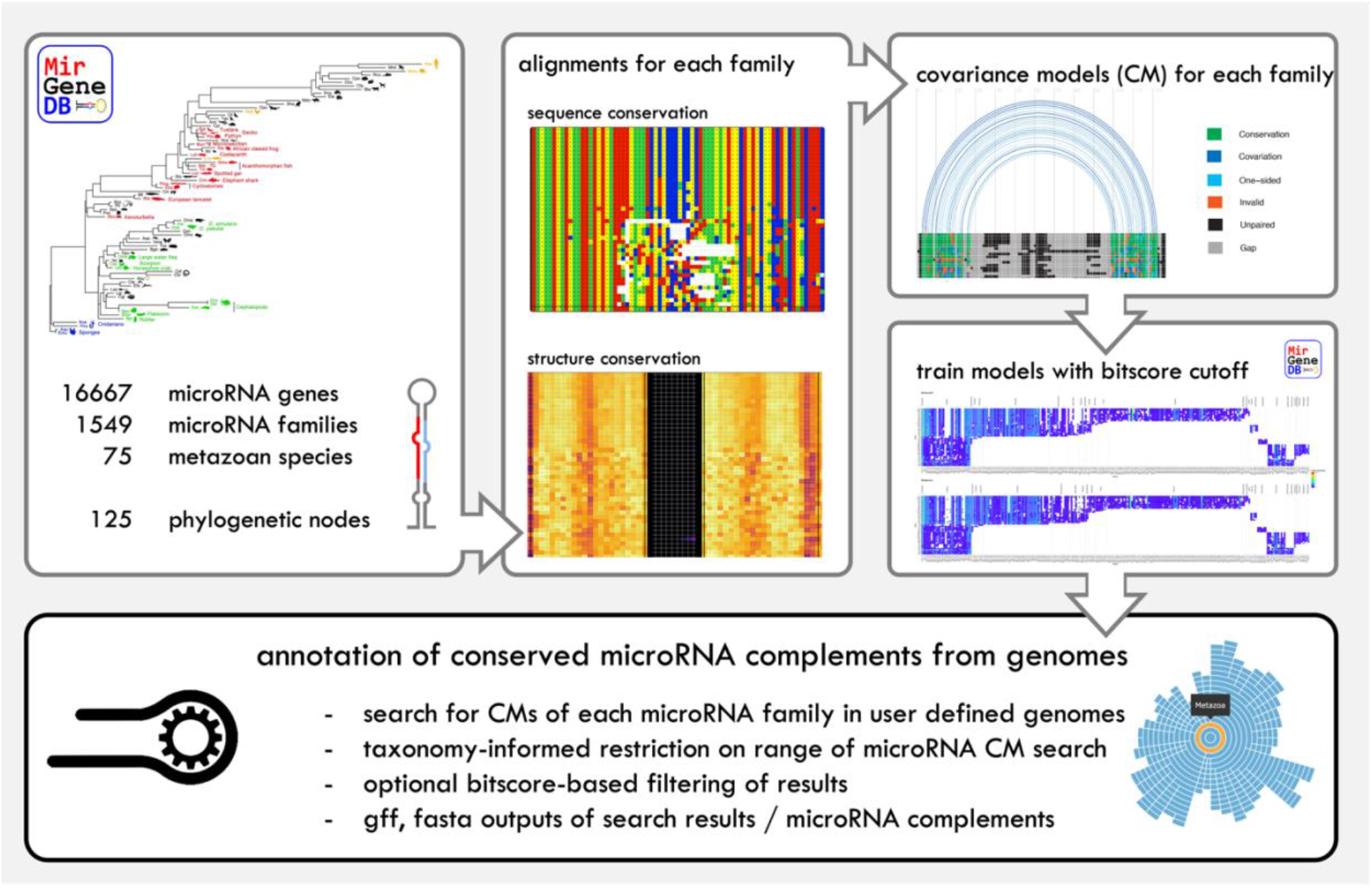

## Introduction

MicroRNAs are among the most conserved regulatory elements in animal genomes and have crucial roles in development and disease^1, 2^. They have long been proposed as disease biomarkers^3–5^, phylogenetic markers for studying animal systematics^6, 7^, and for understanding the evolution of complexity in metazoans^8, 9^. Currently, however, the annotation and naming of *bona fide* microRNA complements requires assembled genome references, small RNA sequencing (smallRNAseq) data from different tissues and developmental stages, and substantial hands-on curation of the outputs from microRNA prediction tools^10–12^. Because these tools were not designed to handle the amount of sequencing data or genome assembly sizes available today and often have high false-positives rates, using them is a tedious process that requires years of training, often extensive computational resources, experience and substantial amounts of time^13^. Especially in larger projects that are not focused on microRNAs, but rather might attempt to annotate them along with other coding and non-coding genes, the required level of attention to detail is often missing which inevitably results in biologically meaningless microRNA results^13–17^, as well as thousands of spurious microRNA annotations^1^. These shortcomings, coupled with the availability of high-quality and publicly available microRNA annotations suited for comparative genomics studies led to the construction of the curated microRNA gene database MirGeneDB^1, 18, 19^. MirGeneDB version 2.1 (2022) now contains microRNA complements for 75 metazoan species spanning all major metazoan phyla over ∼850 million years of animal evolution^19^. Since each gene and family was manually curated in all species in MirGeneDB, highly accurate alignments across this wide span of animal evolution are available that capture a high proportion of the sequence variability for each family. Importantly, each microRNA gene and family is associated with a detailed phylogenetic reconstruction of the evolutionary node of origin and estimated age. This dataset, hence, represents a starting point to better understand features of microRNAs ^20^ and to generate better tools for the prediction of microRNAs. Despite MirGeneDB curating a relatively large number of phyla, the number of species currently covered (75 species) is a far cry relative to the thousands of high-quality animal genomes currently available ^21^ (Figure 1).

**Figure 1:**
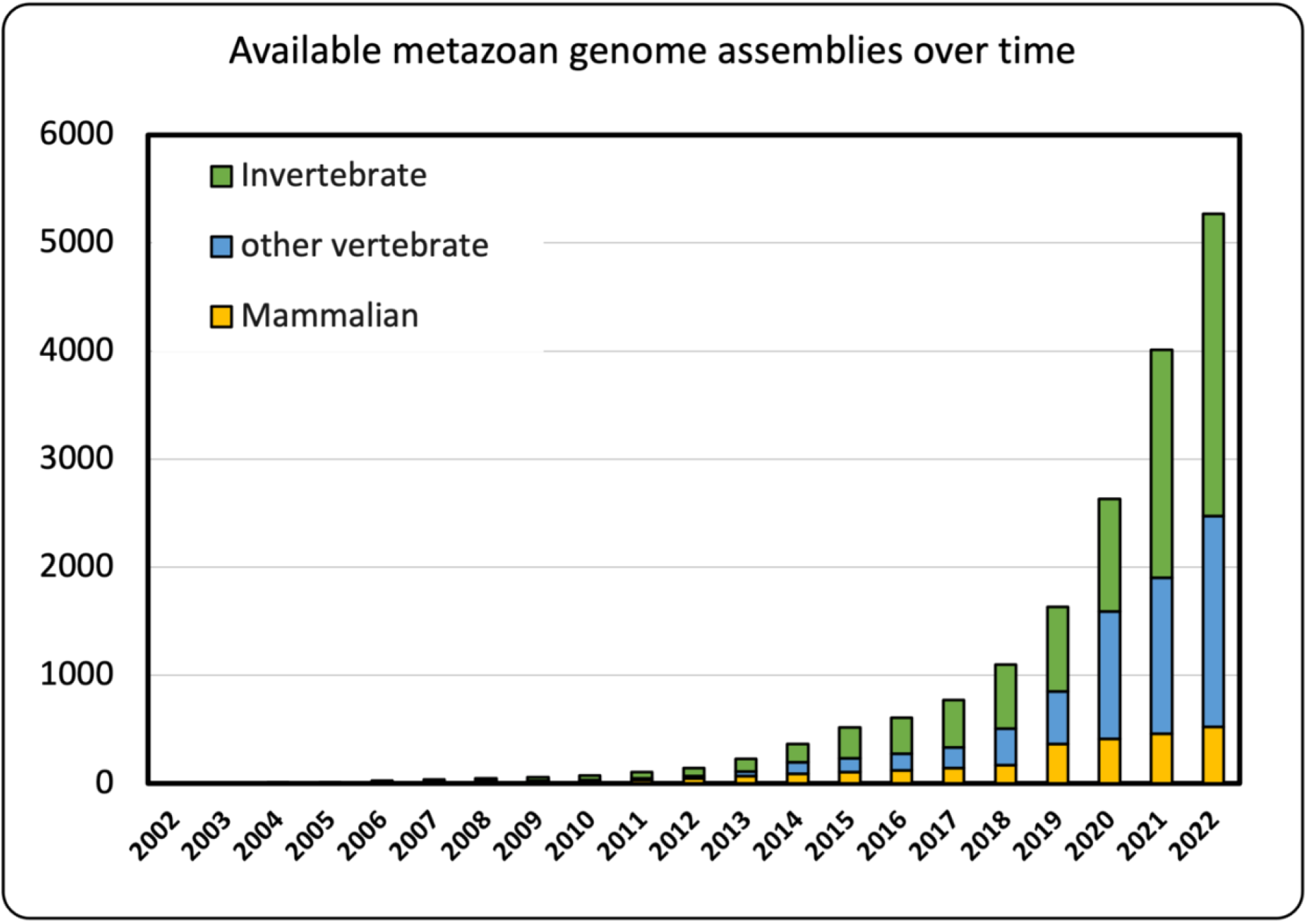
The number of available animal genome assemblies grows exponentially and with more than 5250 currently (2022) available datasets has dramatically grown (Clark et al., 2016).

Very few of these species have been annotated for microRNAs, or have small RNA sequencing data published, thus, comparatively little progress has been made on the suggested microRNA applications (but see ^12, 22–24^ for examples using manual curation). This discrepancy persists because, among other things, no reliable *in silico* method currently exists to annotate conserved or species-specific microRNA complements from genomic references only. Previously, ‘lift-over’ approaches based on whole-genome alignments in model organisms have been used to identify microRNA loci across species ^25, 26^, but it is unclear how accurate these predictions are on the level of the full microRNA complement, or how they computationally scale with size or number of aligned genomes in, for instance, mammals. Despite the availability of computational methods for the search of short RNAs such as microRNAs^27^ and sophisticated machine-learning based tools for non-coding RNA applications^28^, there is currently no approach satisfying the demands of high precision, low false discovery rates and minimized computational demand in a fully automated and user-friendly pipeline^29^. It is a widely acknowledged problem for machine learning applications in genomics in general that existing tools are based on incomplete models^30, 31^. This is the case for microRNA families from miRBase^32^. Such models, for instance, covariance models (CMs) of individual RNA classes, families or genes, as used to group all RNA-families in the Rfam database^32^, are technically quite accurate in detection of many non-coding RNA families^33^. However, these probabilistic models that flexibly describe the secondary structure and primary sequence consensus of an RNA sequence family, require high quality alignments from curated RNAs ideally coupled with detailed evolutionary information to distinguish families and genes over evolutionary time that, until recently, did not exist for microRNAs.

Taking advantage of the manually curated and evolutionarily informed microRNA complements of 75 metazoan organisms in MirGeneDB 2.1^19^, we here built and trained high-quality CMs for 508 conserved microRNA families and integrated them into a fully automated pipeline for microRNA annotation: MirMachine. We show that MirMachine produces highly accurate microRNA annotations in a time-efficient manner from animal genomes of all classes, including very large and recently duplicated genomes, as well as from genomes of extinct species. Using the example of 88 eutherian genomes, we further show that MirMachine predictions can be summarized in a microRNA score that can be used to assess low contiguity or completeness of genome assemblies. MirMachine is freely available (https://github.com/sinanugur/MirMachine) and also implemented as a user-friendly web application (www.mirmachine.org).

## Results

### Accurate Covariance models of 508 conserved microRNA families

16,670 microRNA precursor sequences from 75 species were downloaded from MirGeneDB and all variants from the same genes, antisense loci, and species-specific microRNAs (i.e., not conserved in any other species) were removed arriving at a total of 14,953 genes representing 508 families (Figure 2A).

**Figure 2:**
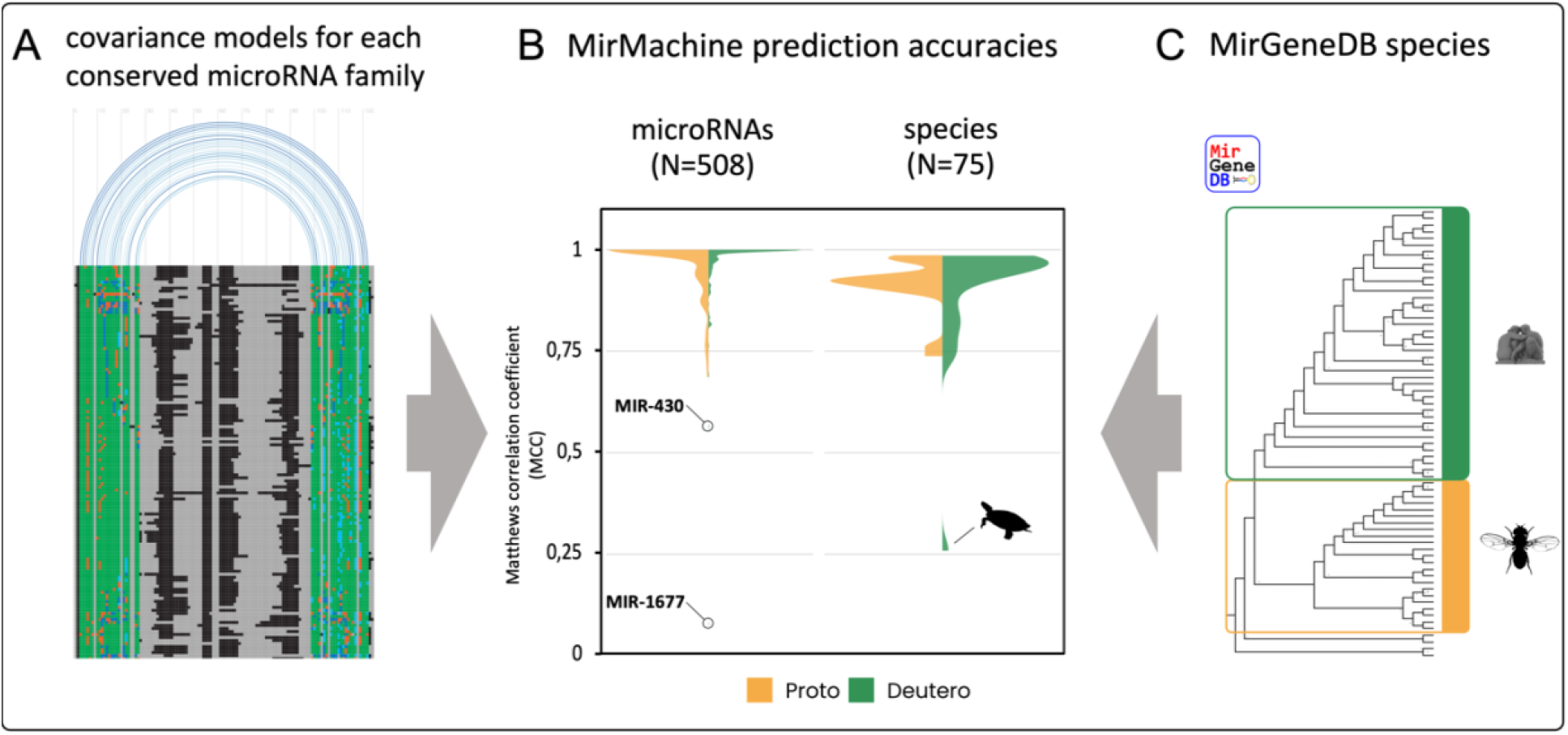
Developing MirMachine covariance models (CMs). A) The MirMachine workflow uses microRNA family-based precursor sequence alignments and structural information to build CMs that B) show very good overall prediction performances when models are run on C) 75 MirGeneDB species using distinct models for protostomes (yellow) and deuterostomes (green) or combined models (not shown).

All microRNA genes for each family were aligned, and covariance models (CM) were built (combined models). Given the evolutionary microRNA family definition used by MirGeneDB, microRNA families can include nucleotide differences in mature and seed that are captured and summarized in the models. To get a finer resolution of our models, we then split deuterostome (N=42) and protostome (N=29) representatives and repeated the process to arrive at 388 microRNA family models for deuterostomes and 143 microRNA family models for protostomes. Depending on the age of a given microRNA family, the number of species that shared the family, the number of existing paralogues and the degree of conservation between orthologues and paralogues, these models contain between very few and many hundreds of individual sequences (see Supplementary Figure 1 for representative examples).

**Supplementary Figure 1:**
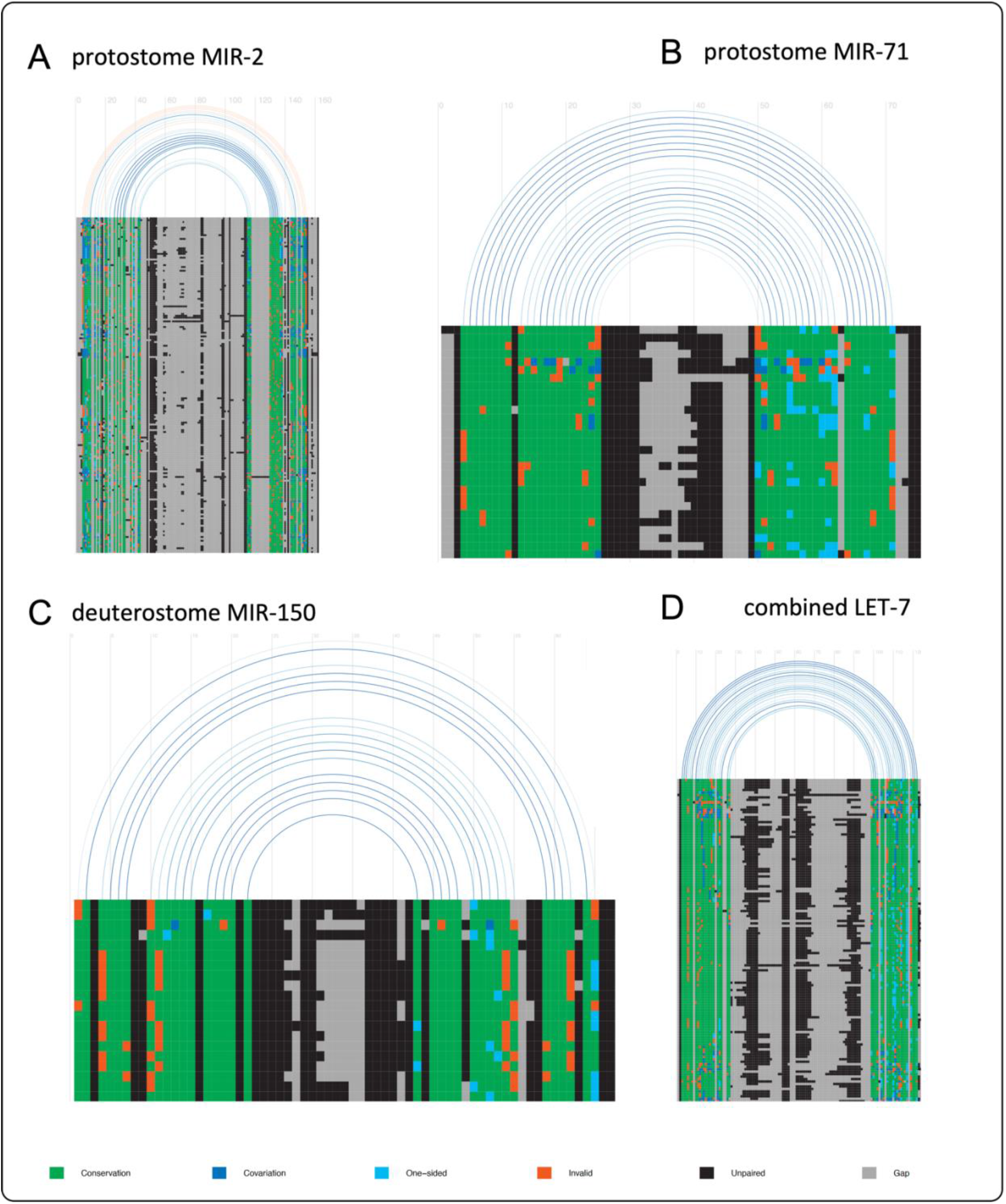
Graphical representation of CMs of representative microRNA families. Conserved base pairs are colored in green. Blue indicates a compensatory mutation relative to the green pairs (dark blue for a double-sided mutation, light blue for a one-sided mutation). Non-canonical paired bases are red, non-base-pairing bases are black. Graphical representations of all CMs used by MirMachine can be found on github (https://github.com/sinanugur/MirMachine-supplementary/tree/main/CM_figures).

Using our workflow (see material and methods), CMs were subsequently trained on the full MirGeneDB dataset to derive optimal cutoffs for their prediction. To measure the prediction accuracy of these models we then used the models on all MirGeneDB species comparing the predictions to the actual complements. An overall very high mean prediction accuracy of 0.975 (Matthews Correlation coefficient (MCC)) for combined models, and 0.975 for deuterostomes, and 0.966 for protostome-models, respectively, was found (Figure 2B, left & Figure 2C). Two microRNA families, MIR-430 and MIR-1677 from the deuterostome models, showed substantially lower MCC scores due to a well-known variability within the MIR-430 family^34–36^ and a combination of low level of complexity and high variation between orthologues in the Diapsida-specific MIR-1677 (Supplementary Figure 2).

**Supplementary Figure 2:**
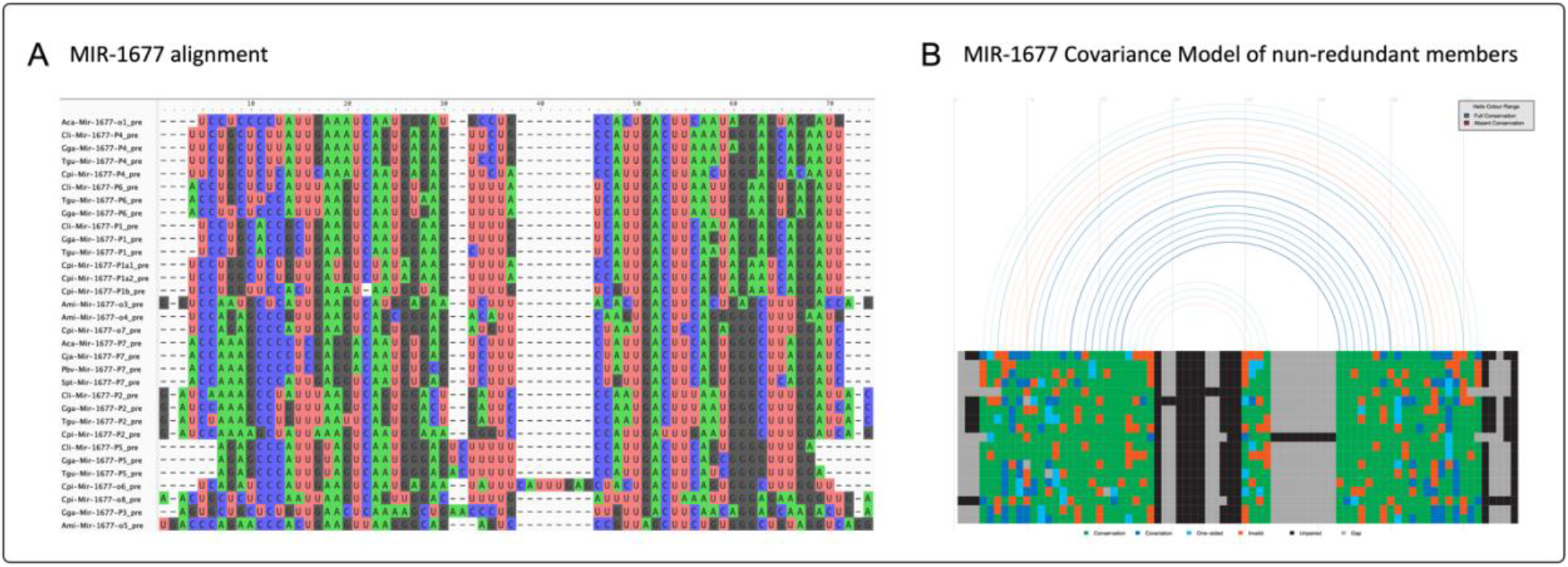
A) Alignment of Mir-1677 genes from MirGeneDB shows low conservation that explains poor performance of B) MIR-1677 CMs in MirMachine.

Conversely, we observe high mean species accuracies of 0.91 for combined models, 0.92 for deuterostomes and 0.92 for the protostome models (Figure 2B, right). The reason that the turtle (*Chrysemys picta bellii*) has such a low MCC is due to the identification of nearly two thousand likely artifactual hits for MIR-1677.

### MirMachine CMs models are largely independent of any single species

To identify potential effects from circular logic of predicting microRNAs of a species that were included to build the query models, we retrained all models for deuterostomes without including human and all protostome models without including the polychaete *Capitella teleta*. Those were chosen because of their relatively complete microRNA complements relative to their respective phylogenetic nodes and given the fact that neither has a sister species in our database (unlike e.g. *Drosophila* or *Caenorhabditis*), which would have heavily biased microRNA recovery. We then used the new deuterostome and protostome CMs to predict microRNA complements in human and *C. teleta*, respectively. We found that MCC for *H. sapiens* only very slightly decreased in accuracy from 0.97 to 0.96 highlighting the robustness of MirMachine covariance models in deuterostomes. In protostomes, the effect on MCC was stronger for leaving out *C. teleta* with a decrease from 0.92 to 0.76. Specifically, some families were not found, including the bilaterian families MIR-193, MIR-210, MIR-242, MIR-278, MIR-281, MIR-375, the protostome families MIR-12, MIR-1993 and the lophotrochozoan family MIR-1994, which were still predicted, but fell below a newly defined threshold. This highlights a markable higher sequence divergence within protostomes, which is likely due to the age of the group, the lower number of representative clades, lower number of paralogues and orthologues per family, and a lower number of species in general. The annelid families MIR-1987, MIR-1995, MIR-2000, MIR-2685, MIR-2687, MIR-2689 and MIR-2705 were not searched because no models were built given the absence of a second annelid species, highlighting the importance of including at least two representative species for each clade in MirGeneDB^19^.

### Performance of MirMachine prediction versus MirGeneDB complement

To get a comprehensive understanding of the performance of MirMachine on the microRNA complements of MirGeneDB species, we looked in more detail at the performance of CMs, and their respective cut-offs, for a selection of major microRNA families (N=305) including all gene-copies (N=12,430) (Figure 3). When comparing the MirGeneDB complements (Figure 3A) with the predictions from MirMachine (Figure 3B), similarities were striking and overall differences limited to few families (Figure 3C); indicating either potentially false positives (231) or false negatives (421), respectively (Supplementary File 1). These are of further interest as they either represent missed microRNAs in MirGeneDB, or significant deviations from the general CMs and, hence, possibly incorrectly assigned microRNA paralogues in MirGeneDB.

**Figure 3:**
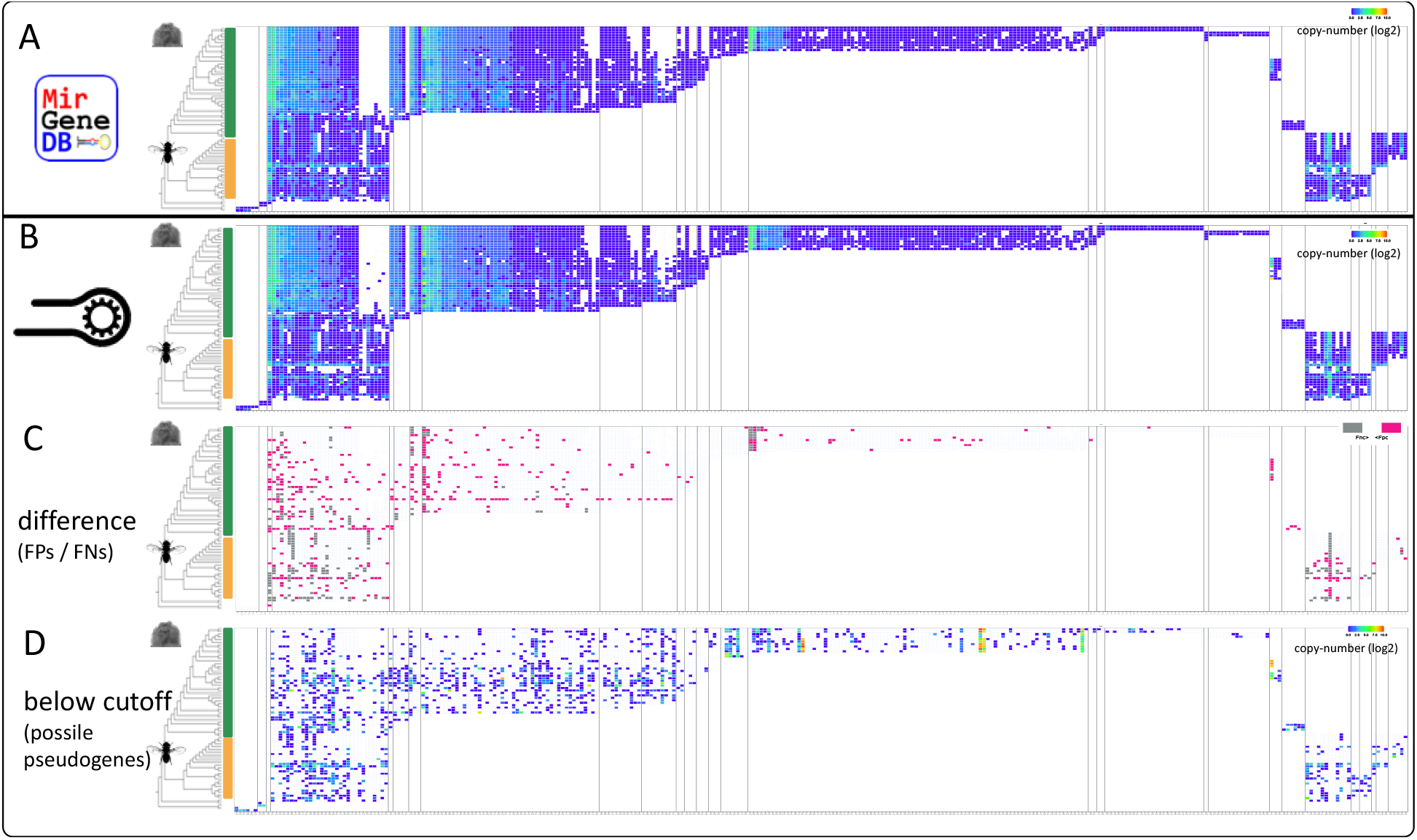
Detailed comparison of MirMachine predictions on 75 MirGeneDB species and 305 representative microRNA families in the form of banner-plots. Columns are microRNA families sorted by phylogenetic origin and rows are species. Heatmap indicates number of paralogues / orthologues per family. A) the currently annotated microRNA complements in MirGeneDB 2.1^19^. B) MirMachine predictions for the same species and families show very high similarity to A. C) Differences between A and B highlighted as potential false-positives (pink) or false negatives (gray). D) MirMachine predictions below cut-off based on training of CMs on MirGeneDB show a range of potential random predictions and pseudogenes, highlighting the effect of curation & machine learning on models.

Finally, we found a substantial number of low-scoring MirMachine predictions of microRNA families that did not reach the determined cutoff based on trained CMs (Figure 3D) and therefore are not considered *bona fide* microRNAs. However, we found that these also contain pseudogenized microRNA orthologues (or paralogues) exemplified by a hitherto unknown human LET-7 pseudogene that is not found expressed in any MirGeneDB sample (Figure 4). To our knowledge, this is the first report of, and MirMachine the respective tool for, pseudogene-predictions of microRNAs. Pseudogenes, or ‘gene-fossils’, are potentially very useful to determine the rate of gene duplication and follow the evolution of sequence changes in organisms and might be included in studies studying cause and consequences of duplications on microRNAs^23^.

**Figure 4:**
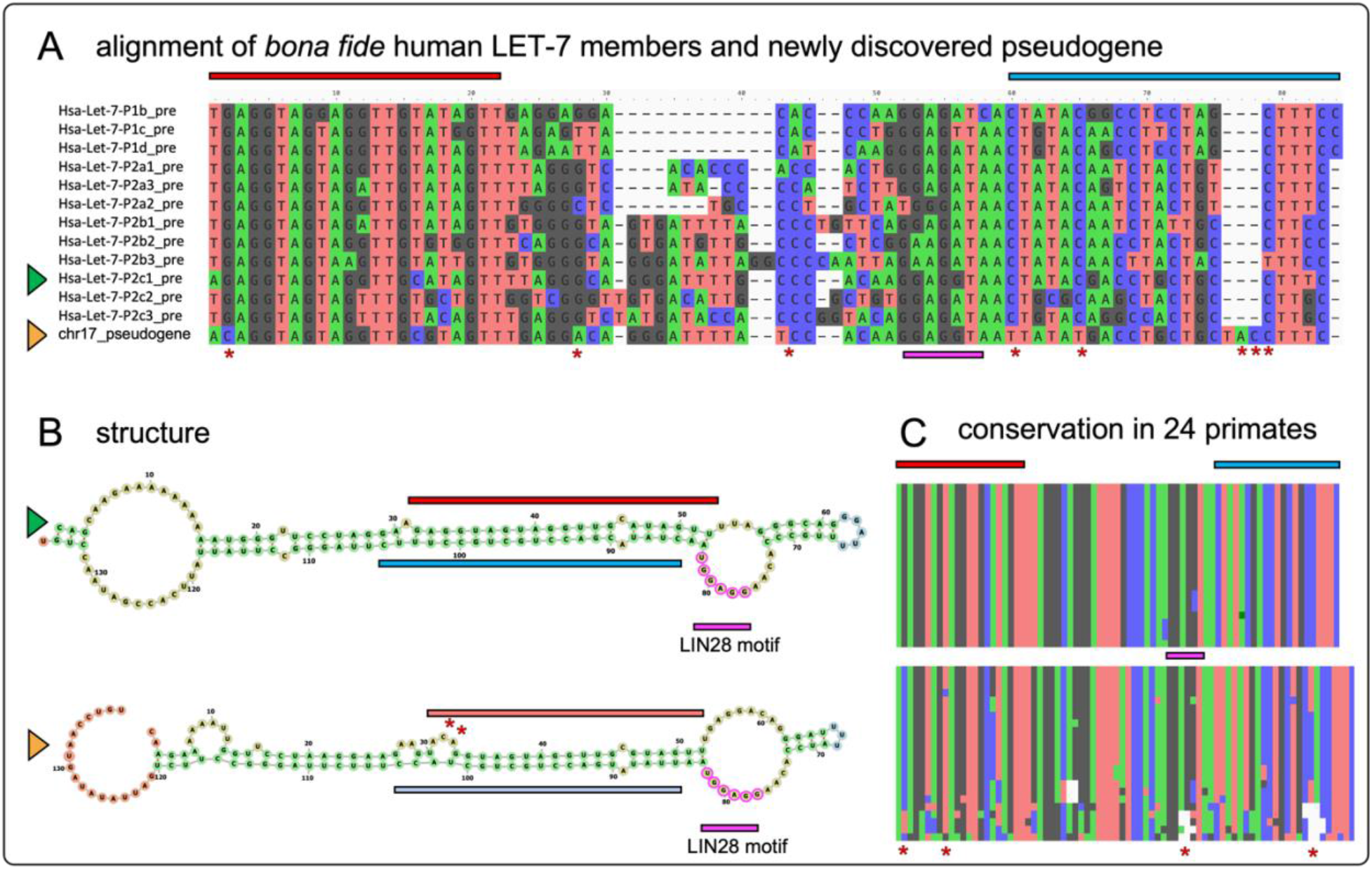
The human Chr.17 LET-7 pseudogene. A) sequence alignment of the currently annotated 12 *bona fide* LET-7 family members in human and the pseudogene candidate discovered by MirMachine. Non-random sequence similarities, including LIN28 binding sites (pink) are apparent with few noteworthy differences (asterisks) such as in position 2 on the 5’ end (red box indicates mature annotation, position 2 equals seed-sequence) or a triplet insertion at the 3’ end (blue box indicates star sequence annotation) are indications for non-functionality. B) Structural comparison of a representative *bona fide* LET-7 member (Hsa-Let-7-P2c1, green triangle) with the pseudogene (yellow triangle) highlights similarities of pseudogene candidate to *bona fide* microRNA, but points out disruptive nature of nucleotide changes for the structure (asterisks) very likely affecting a potential Drosha processing. C) sequence conservation of *bona fide* Hsa-Let-7-P2c1 (top) and the pseudogene (bottom) in 24 primate genome (ENSEMBL v100) highlights the sequence conservation of *bona fide* microRNAs from the loop showing some changes, the star (blue) few changes and the mature (red) showing none, while the pseudogene shows many more changes and seems to be enriched in disruptive changes in the mature / seed region.

### The microRNA complements of eutherians reveal the microRNA score as simple feature for genome contiguity

Applying MirMachine to a testcase, we downloaded 89 eutherian genomes currently available in Ensembl that are not curated in MirGeneDB and annotated their conserved microRNA complements. Altogether 38,550 genes in 260 families, in about 4,400 CPU hours, were found and showed an overall very high concordance between species (Figure 5A). As expected, Catharrini (pink) and Muridae (light green) specific microRNAs were only found in the respective representatives, but surprisingly, six species (Figure 5, yellow arrows) showed substantial absences of microRNA families. We therefore wondered whether these absences indicate microRNA losses due to biological simplifications (see ^22^), proposed random events^37, 38^, or whether they might be due to technical reasons^7^. Given that the outlier species (Alpaca, Shrew, Hedgehog, Tree shrew, Pika, and Sloth) have no particularly reduced morphology, we reasoned that the source might be technical and recovered N50 contiguity values for all genomes. We found that all six genomes had substantially lower N50 values than all other genomes, indicating that microRNAs might be able to predict completeness of genome assemblies (Figure 5B). Therefore, we next developed a simple microRNA scoring system defined as the percentage of expected conserved microRNA families found from a genome (in this case including 175 microRNA families found in most eutherians according to MirGeneDB ^19^, and showed that microRNA scores below 80% correlate with very poor N50 values <10kb and that N50 values of 100kb indicate microRNA scores of 90% and higher (Figure 5C, red and blue lines). A noteworthy exception is the microbat *Myotis lucifugus* with a N50 of 64kb and a microRNA score of 74%, which might be explainable by previously suggested genome evolution mode through loss^39, 40^.

**Figure 5:**
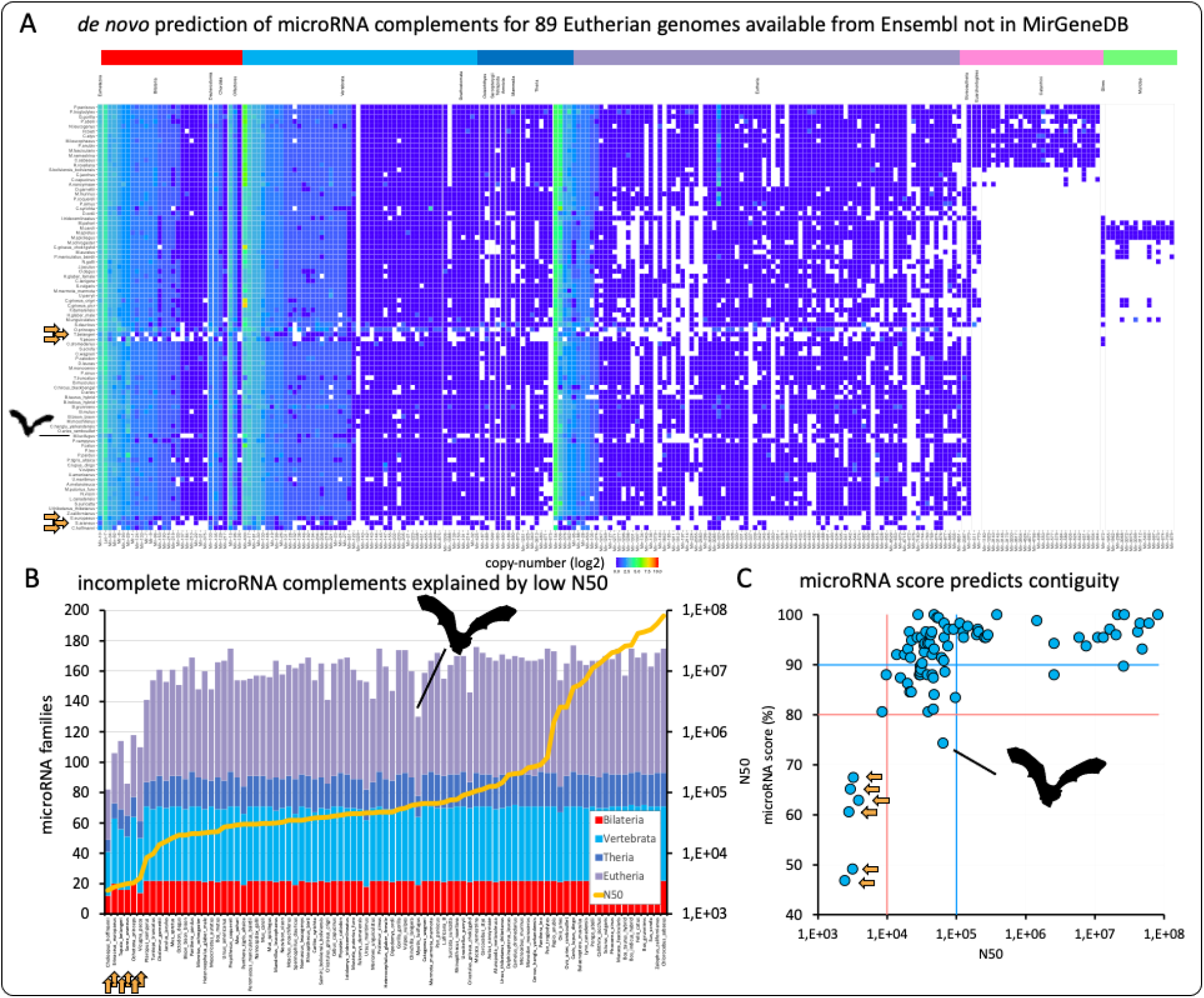
MirMachine predicts conserved microRNA complements of 89 eutherian mammals available on Ensembl and not currently represented in MirGeneDB. A) banner plot of results for MirMachine predictions on 88 eutherian mammalian species for selected range of major microRNA families and genes showed very strong homogeneity of microRNA complements in general and identified a number of clear outliers (yellow arrows, including Alpaca, Shrew, Hedgehog, Tree shrew, Pika, and Sloth). B) Stacked histogram sorted by N50 values). Outlier species (yellow arrows: same as in A)) all have very low N50 values, indicating an artificial absence of these phylogenetically expected microRNA families. C) The microRNA score predicts the assembly contingency and is the proportion of phylogenetically expected microRNA families that are found in respective genomes (here eutherians). microRNA scores below 80% (red horizontal line) tend to have low N50 values (red vertical line indicates N50 below 10,000 nucleotides), while scores above 90% indicate N50 higher than 10,000 nucleotides. Noteworthy exception is the bat *Myotis lucifugus* which might be explainable by previously suggested genome evolution mode through loss 39,40.

### MirMachine predicts microRNAs from extinct organisms and very large genomes

High quality *in silico* annotation of genomes is particularly important for organisms where no high quality RNA is likely to ever become available. This is the case for species such as mammoths that went extinct millennia or even millions of years ago (but see ^41^). Using available data from extinct and extant elephantids^42, 43^, we ran MirMachine on 16 afrotherian genomes, including the hyrax (*Procavia capensis*) from Ensembl and the tenrec (*Echinops telfairi*) from MirGeneDB, and 14 elephantids including extant savanna elephants (*Loxodonta africana*), forest elephants (*Loxodonta cyclotis*) and asian elephants (*Elephas maximus*) respectively (Figure 6A, green elephantid silhouettes), but also extinct american mastodon (*Mammuthus americanum*), straight-tusked elephants (*Palaeoloxodon antiquus*), columbian mammoth (*Mammuthus columbi*) and the woolly mammoths (*Mammuthus primigenius*) (Figure 6A, red elephantid silhouettes). We find a very high degree of similarities between afrotherians, and striking congruence between extinct and extant species which indicates the high accuracy of the MirMachine workflow. More so we find patterns of microRNA losses that could be phylogenetically informative (Figure 6A, arrows). For instance, we do not find MIR-210 in any of the elephant species, which might be a elephantid specific loss (Figure 6A, pink arrow), we further find that *P. antiquus* and *L. cyclotis* have both lost MIR-1251 (Figure 6A, light blue arrow), and a shared loss of MIR-675 and MIR-1343 (Figure 6A, purple arrows), both supporting previously identified sister group relationships^42^.

**Figure 6:**
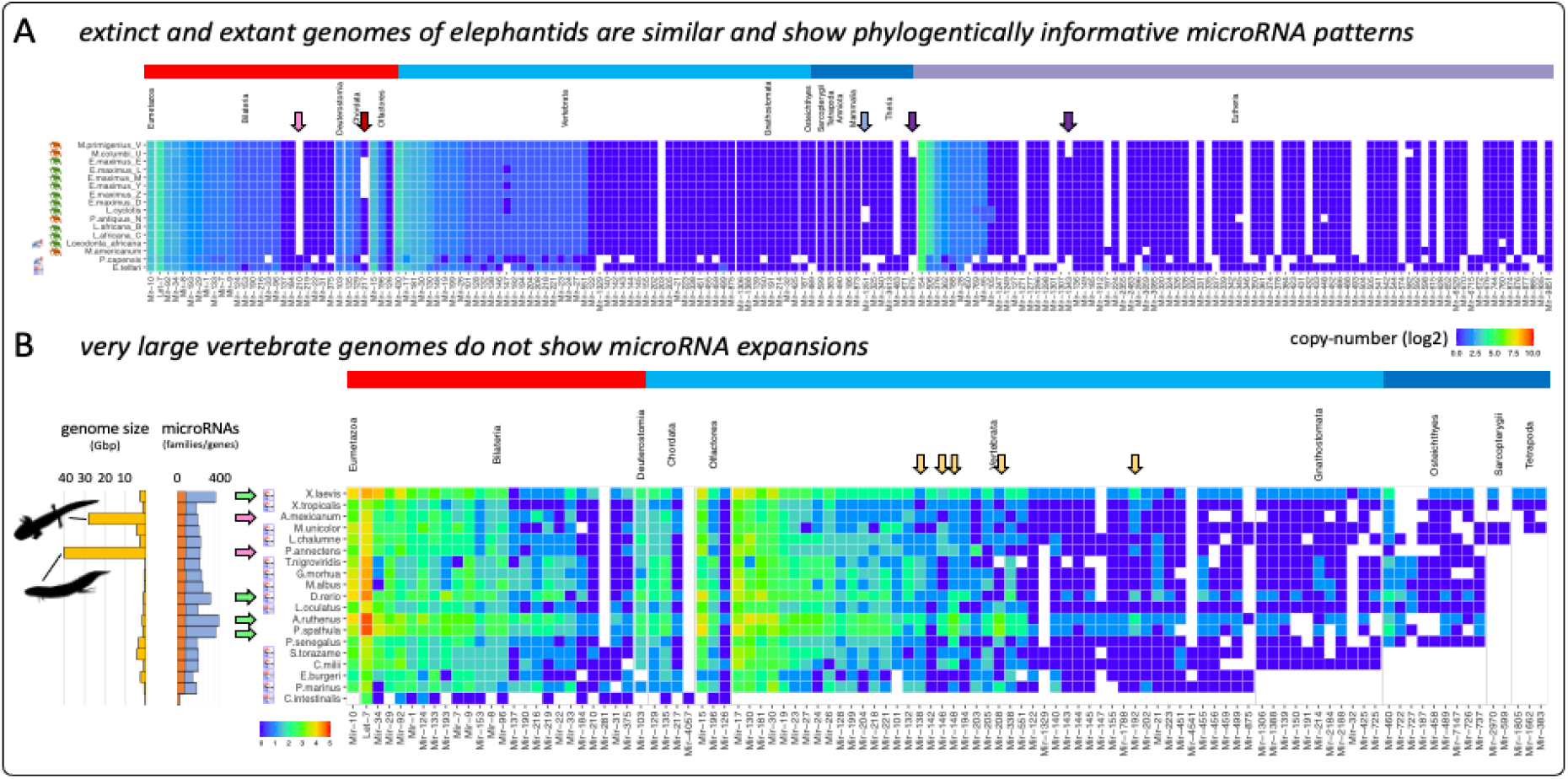
MirMachine enables microRNA complement annotations from extinct and very large genomes. A) MirMachine predictions from afrotherians show no clear differences between extinct and extant genomes, but likely phylogenetically informative losses of microRNA families (colored arrows). B) MirMachine predictions in organisms with extensive genome expansions (pink arrows) show no expansion of microRNAs, but organisms with known genome duplications (green arrows) do. A number of shared microRNA copiews in sterlet (*A. ruthenus*) and paddlefish (*P. spatula*) support a common genome duplication event in the last common ancestor of Acipenseriformes (yellow arrows).

A pertaining challenge for microRNA prediction and annotation of extant species, is the occurrence of additional whole genome duplication events and, not necessarily connected, extreme genome expansions. This often leads to computational challenges where identical copies are hard to distinguish based on read-mappings or genomes are simply so large that existing pipelines need extensive computational resources often facing programmatic limits. Therefore, we next investigated the performance of MirMachine in vertebrate species with very large genomes and of known additional rounds of genome duplications. For the first group, we included the axolotl (*Ambystoma mexicanum*) with a genome of 28 Gbp and the african lungfish (*Protopterus annectens*) with a genome of bigger than 40 Gbp into our analysis. For the second group we included the African clawed frosh (*Xenopus laevis*) with known allotetraploid genome^44^ and the zebrafish (*Danio rerio*) from MirGeneDB, the sterlet (*Acipenser ruthenus*) with proposed sturgeon specific genome duplication and occurrence of segmental rediploidization^45^, as well as the american paddlefish (*Polyodon spathula*) with a recently shown genome duplication which was, however, interpreted as sturgeon independent^46^. We combined these species with the gray bichir (*Polypterus senegalus*) that has a moderately sized (e.g., human-sized) genome and no unique known genome duplication events, along with 13 other MirGeneDB species representing a range of Olfactores, vertebrates, gnathostomes, Osteichthyes, Sarcopterygii and Tetrapoda representatives (Figure 6B). We find that MirMachine ran very well on all genomes using 32 cores and under 2 hours per species, whereas the lungfish ran the longest (around 3 hours 45 mins). As expected, we find that the size of the genomes do not affect the microRNA complements (Figure 6B, pink arrows), but that organisms with additional whole genome duplications (Figure 6B, green arrows) clear trace of duplications (also see ^23^). A curious observation was that sterlet and paddlefish showed very consistent microRNA copy-number patterns, in particular in the retention of additional MIR-138, MIR-146, MIR-148, MIR-192 and MIR-208 copies (Figure 6B, orange arrows) indicating a likely common origin of genome duplication at the last common ancestor (Acipenseriformes), or very similar retention pressure in the more unlikely case of independent duplication. Altogether MirMachine is a suitable tool for the annotation of microRNA complements from extinct and very large genomes alike.

### MirMachine models outperform existing Rfam models

In the most recent Rfam update (v. 14) an expanded assembly of microRNA models based on miRBase was released^32^. As mentioned here before, and stated elsewhere, a major concern in microRNA research has been the quality of this online repository of published microRNA candidates^1, 47–58^ with estimates of two out of three false-positive entries. Thus, the database contains more false positives than microRNAs. These are for instance numerous tRNA, rRNA or other fragments, but also incorrectly annotated *bona fide* microRNAs that strongly influence interpretations of data. In addition to the false positives, numerous miRBase annotations are imprecise and have varying precursor annotation forms (with or without flanking regions of varying lengths) and not both arms are annotated, 3’ ends are incorrect, and in a few cases even 5’ are not correctly annotated which substantially affects target predictions (for details see ^1^). Further, it uses an outdated nomenclature which is inconsistent in that members of the same microRNA family are not named the same way making the identification of family members cumbersome. This problem has to a large extent been transferred to Rfam and their microRNA family models in particular (e.g. MIR-95 family member Hsa-Mir-95-P4 (https://mirgenedb.org/show/hsa/Mir-95-P4) with own model https://rnacentral.org/rna/URS0002313758/9606, or MIR-15 member Hsa-Mir-15-P1d https://mirgenedb.org/show/hsa/Mir-15-P1d) with own model (https://rnacentral.org/rna/URS000062BB4A/9606 (see Supplementary File 2). This all has been addressed in the manually curated microRNA gene database MirGeneDB.org^1, 19^ and MirMachine, respectively.

Regardless, we tested the performance of 523 Rfam microRNA models, that we curated to be of animal origin, on the 75 MirGeneDB species and found that 36,931 microRNAs were predicted (compared to 16,913 MirMachine and the 15,846 microRNA annotations in MGDB 2.1). Given that the number of conserved microRNA families is a focus of MirGeneDB and very unlikely to be expanded in the future^13^, this much higher number of predictions suggests that Rfam predictions contain thousands of false positives (FPs). We further looked for performance of highly conserved families (see materials and methods). Rfam models had MCCs of 0.96, 0.94, 0.96 and 0.89 for microRNA families LET-7, MIR-1, MIR-196 and MIR-71 respectively. The same family performances for MirMachine were 0.97, 0.98, 0.97, 0.97. Thus, as expected, Rfam model had comparable performance for these correctly assigned, and deeply conserved families, but performed poorly for incorrectly assigned microRNAs.

### MirMachine outperforms whole genome alignment approaches

We compared the performance of MirMachine with a whole-genome alignment approach as used previously in ‘lift-over’ approaches in e.g. *Drosophila* genus ^25, 26^. Using the 470-way mammalian species MULTIZ genome alignment based on the human genome, we tested how accurate these predictions are on the level of the full microRNA complement and how they computationally scale with size or number of aligned genomes in, in this case, mammals. We find that most human microRNA loci indeed produced alignments in most species, but that there was a substantial number of 1) missing families and genes and 2) a very high number of false positives calls in these microRNA alignments (Supplementary Figure 3 & github).

**Supplementary Figure 3:**
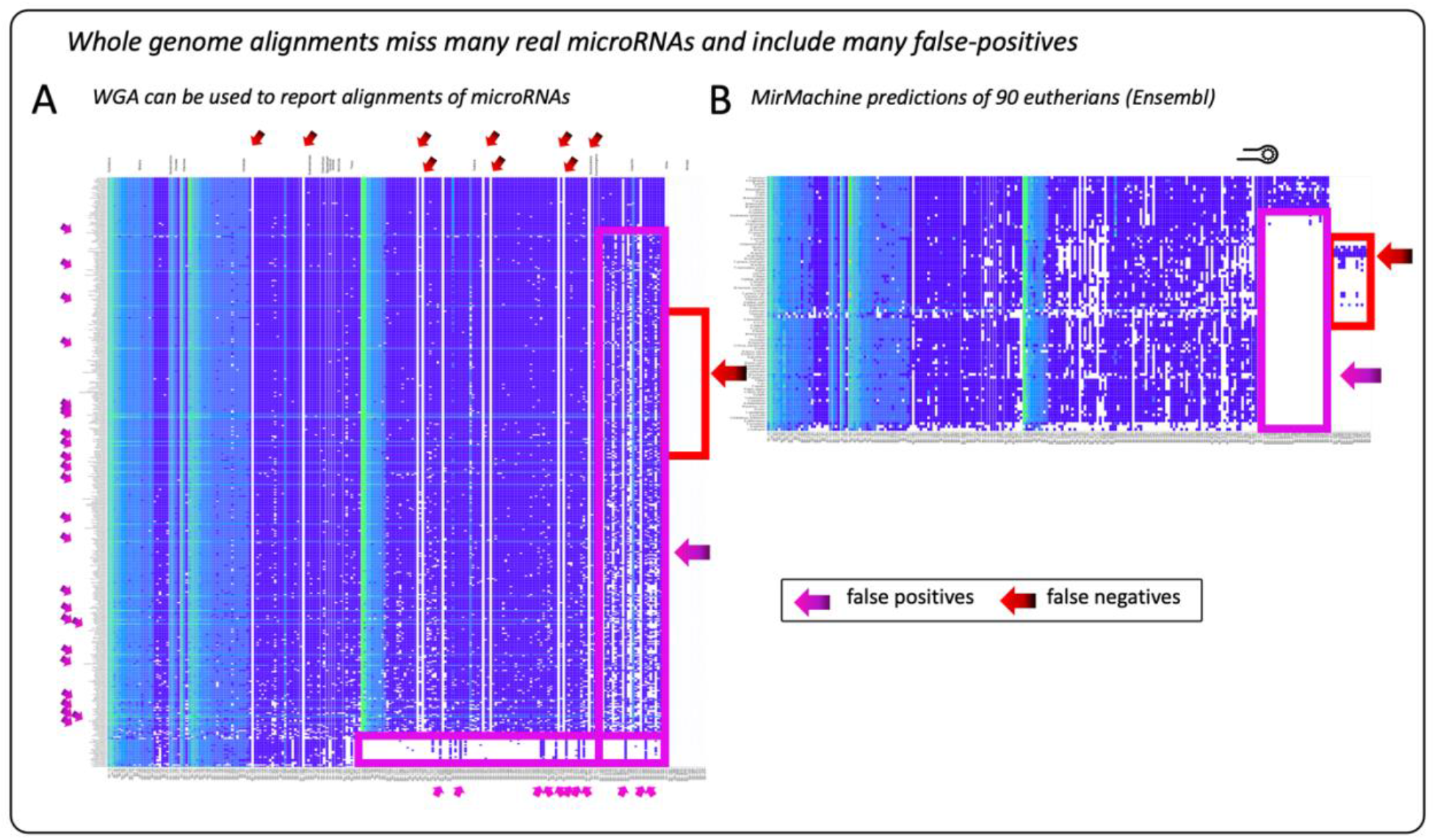
When comparing overall performance of (A) alignments reported for each of the 470 mammalian species, the overall impression is that many microRNA loci in human are aligned in a majority of mammalian genomes. However, when comparing to the MirMachine output (B), a number of *bona fide* microRNA families are not reported (red arrows) due to their absence in the human reference (red box: murid microRNA families). Additionally, a high number families and genes that are not expected (pink boxes) given the phylogenetic level of the species (i.e. not Eutherian, not Catharrini) is reported, which seems unlikely to be correct. This also goes for very high number of copies in a number of species (pink arrows left site of A) that would indicate genome duplication, which have not been reported, and likely are false calls.

Specifically, on average, for the 90 eutherian genomes we had previously analyzed with MirMachine, more than 90 false positives per species were reported from WGA on average (Supplementary Figure 4).

**Supplementary Figure 4:**
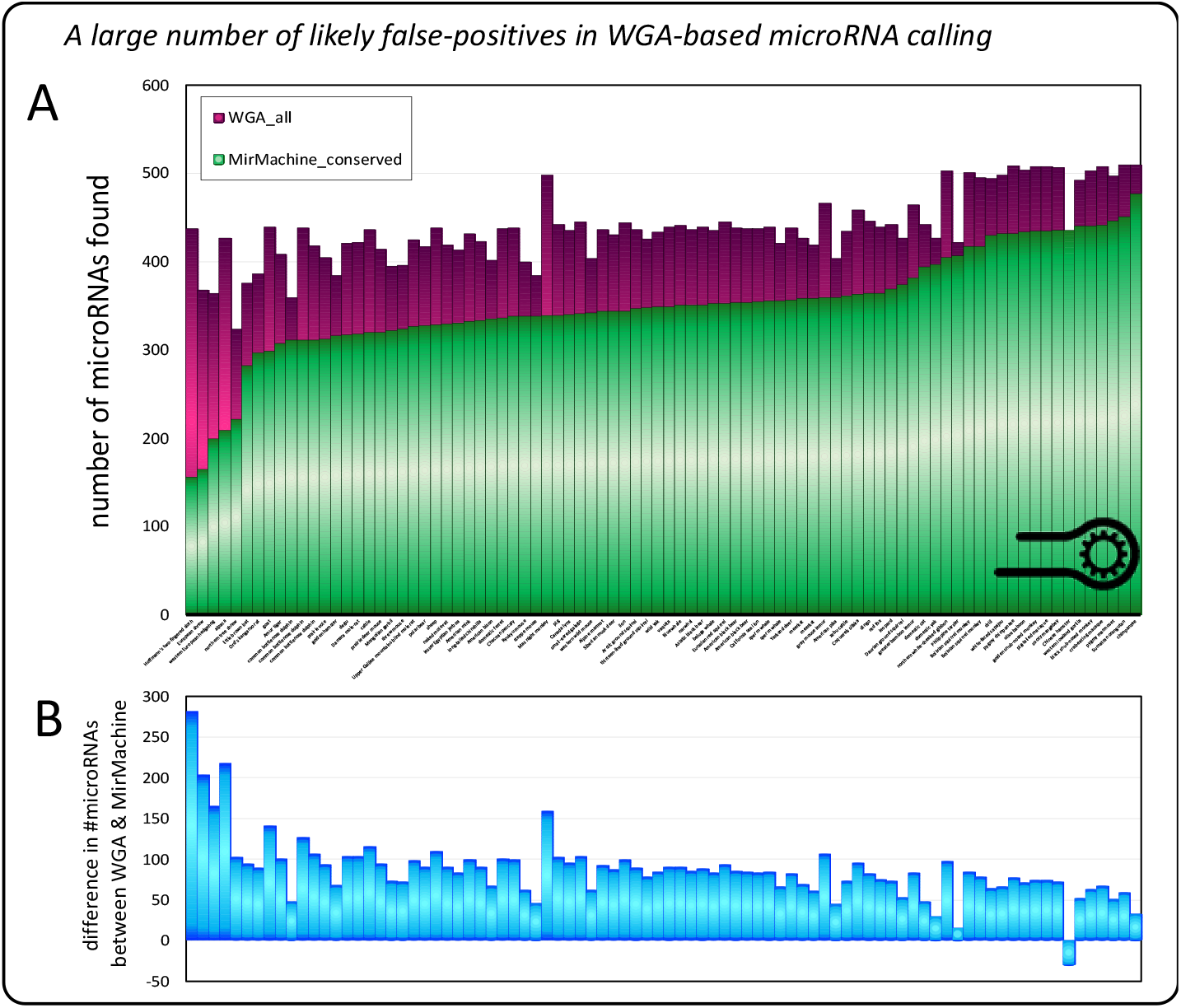
Comparison of the subset of species from the 470 MULTIZ WGA (A – pink) and our Ensembl based 90 eutherians analysis (A –green). On average, more than 90 false positives are found per genome using WGA (B).

To investigate the nature of these likely false calls, we selected 10 microRNA families (see Supplementary Figure 3 small pink arrows at the bottom) with origin in eutherians and Catharrini that were reported in non-eutherians and outside Catharrini, respectively, and carefully checked all alignments to investigate sequence conservation (Supplementary Figure 5). We found that alignments reported from outside the expected groups are too distinct from the reference and are obviously no microRNAs. In an attempt to verify the effect of nucleotide difference between bona fide genes and the aligned regions bearing substantial changes, we took the example of Catharrini-specific MIR-4677 (Supplementary Figure 5 B&C) and, for subset of representative mammals, made structure predictions and were able to show that already slightly changed locus in other primates created structures less likely to be processed as microRNAs (middle structure), with other non-primate mammals showing almost random structures (yellow bar). These results clearly show that WGA based approaches have pitfalls that the MirMachine pipeline avoids.

**Supplementary Figure 5:**
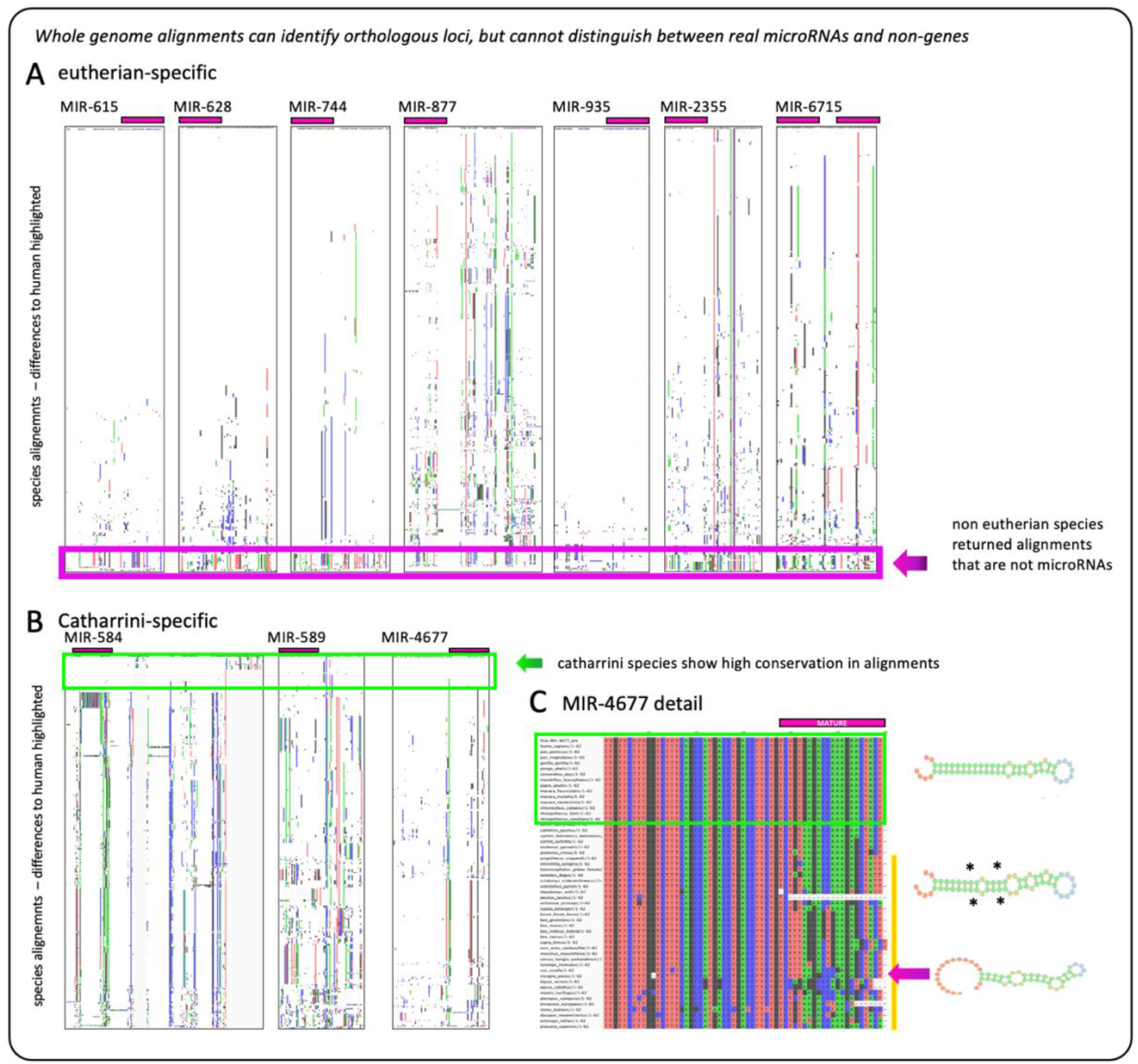
Unexpected microRNA alignments show substantial variation in species not belonging to the group of known evolutionary origin of the microRNA family. Selection of unexpected reports of microRNA presence of A) eutherian-specific and B) Catharrini-specific microRNA families in non-eutherian and non-Catharrini species shows that, while having alignment reported, substantial difference in their aligned-to sequences. This indicates that these are either 1) incorrect alignments or 2) that aligned loci do not contain microRNA genes. In C) (MIR-4677 detail) clear differences in nucleotide composition shows the effect of these sequences on the actual structure of the putative microRNAs clearly ruling out a processing as microRNA. A&B) Each plot highlights the differences to the human reference (white = 100% conserved sites)

### MirMachine functions & options

All models (total, protostome and deuterostome) were implemented into the standalone MirMachine workflow which is available under https://github.com/sinanugur/MirMachine, and the web app www.mirmachine.org. MirMachine also contains the curated “node of origin” information from MirGeneDB that can be used to limit the microRNA gene search to phylogenetically expected microRNA families, substantially reducing the search space and shortening the necessary run-time. Several other options, such as the search for single families (e.g. “LET-7”) or families of a particular node (e.g. “Bilateria”) are available, too. In the web app, genome accession numbers can be provided avoiding the need for down- and upload circles.

## Discussion

The existence of thousands of animal genome assemblies is massively mismatched by the availability of annotations of important gene-regulatory elements such as microRNAs. Here, we have presented MirMachine as an important first step to overcome this discrepancy and the need for small RNA sequencing data or extensive expert manual curation. This is particularly valuable for organisms, tissues, or developmental time points, where expression datasets will be very difficult to acquire and, hence, microRNA detection based on smallRNA sequencing reads impossible. The unique combination of well-established covariance model approaches trained on manually curated and phylogenetically informed microRNA family models built from more than 16,000 microRNAs of 75 metazoan species makes MirMachine very sensitive to detect paralogues of a family in a given organism (low false-negative rate) and very robust against wrong predictions (low false-positive rates). MirMachine’s ability to accurately predict full conserved microRNA complements from genome assemblies, as exemplified by our analysis of nearly 90 eutherian genomes from Ensembl, will not only enable large comparative microRNA studies and automated genome annotation for microRNAs, but also showed the potential of microRNAs for the assessment of genome assembly completeness (Figure 5). Because of the near-hierarchical evolution of microRNAs, they have a very strong potential not only as taxonomic markers as used in e.g. miRTrace^59^ or sRNAbench^60^, but to also outperform approaches that are based on selected sets of protein-coding genes such as BUSCO^61^ or OMArk^62^ (Paynter et al in prep). Those heavily rely on the correct identification of orthologues of selected single copy protein-coding genes, which are much more variable than microRNAs, do only represent a subset of protein coding genes and, hence, cannot be used to accurately assess or measure rates of genomic loss or completeness directly. By comparing N50 values and a herein established microRNA score, we have shown that microRNA complements predicted by MirMachine are suited to assess genome completeness and contiguity. This might have wide-reaching consequences for future applications as a microRNA score could become a standard measure for genome annotation pipelines.

We have also shown that it is possible to use MirMachine’s ‘below cutoff’ predictions for the study of pseudogenes, which could enable better understanding of dosage-level regulation or gene- and genome duplication events, in general^23^. Using several so far uncharted vertebrate genomes of either extreme size (axolotl, lungfish) and comparing them to smaller, but secondarily duplicated genomes, we could show that MirMachine works on such large genomes and confirm that the size of assemblies does not matter for the number of microRNAs, but that genome duplication events do. By directly comparing the outputs of MirMachine counts for microRNA paralogues in sterlet and paddlefish, we found patterns of microRNA duplicates that support a common genome duplication of the two species.

Finally, we employed MirMachine on extinct species genomes’ and could show that besides similarity to extant representatives, several absences / losses of microRNAs were observed within the elephantids that suggest a phylogenetic signal. These findings are exciting as they might give clues on the genome regulation differences in organisms, where actual RNA will be hard or impossible to get by. Importantly, at this stage, we have not yet made sequence-based comparisons of the microRNAs between any of the species. This is an untapped area for future development.

A comparison to whole genome alignment (WGA) approaches revealed that there is indeed a high number of alignments in mammalian genomes relative to human microRNA loci, but that there are several false positive and false negative calls rendering this approach as inferior to MirMachine. However, the identification of loci that do show sequence similarity, but have no microRNA function could be an interesting avenue for future research on the evolution and pseudogenization of microRNAs. Furthermore, WGA based approaches aiming at microRNA complement wide analyses require substantial computational resources and skills and, hence, should not be considered sustainable for the standardized annotation of full microRNA complements.

MirMachine currently provides predictions as community standard file formats GFF or FASTA that are named by family and coordinates, but not according to their possible paralogue or orthologue nomenclature^1^. This is due to the fact that the required syntenic information is often not available and not currently analyzed by our pipeline. Furthermore, MirMachine does not predict species specific microRNAs which can play crucial roles in evolution^24^. MirMachine predictions are a solid foundation for future smallRNAseq driven annotation efforts of novel microRNAs and synteny-supported annotation of paralogues and orthologues.

Per design MirMachine can only predict conserved microRNAs based on MirGeneDB-derived CMs. However, there are a number of tools to predict novel microRNA candidates from genomes using different methodologies but are all not based on a curated reference and, hence, might be of limited value (see ^63, 64^). We strive to address those issues in the future, but would like to stress, in the meantime and in general, that manual curation is a crucial step that should never be disregarded, even though MirMachine heavily reduces the need for extensive and week-long efforts.

The decision to create protostome and deuterostome specific microRNA family models can be seen as a first step toward group-specific microRNA gene-family models that might increase the accuracy of MirMachine further in the future. Variability of model performance based on evolutionary age of families has not been studied here, but the addition of more taxa to MirGeneDB will be an invaluable improvement for group-specific microRNA family prediction and paralogue-specific modeling of microRNAs. We stress that for pre-bilaterian groups of Cnidaria and Porifera MirMachine currently only provides a small set of microRNA models, as these groups show comparable little conservation of their microRNA complements and aberrant microRNA structures^65–68^. Another important area of possible expansion clearly are plant microRNAs, that currently suffer from multiple non-overlapping available databases and potentially stronger curation problems than observed in animals (see ^58, 69^).

MirMachine is freely available as a standalone tool or web application. It enables even non-microRNA experts to annotate conserved microRNA complements regardless of the availability of small RNA sequencing data. Thus, it has a strong potential to close the ever-increasing gap between existing high-quality genomes^70, 71^ and their microRNA annotations. A possible addition of MirMachine into the standard genome annotation pipelines of Refseq and Ensembl is currently discussed. The availability of thousands of metazoan genomes and their microRNA annotations will pave the way toward the promise of microRNAs and a true postgenomic era.

## STAR★Methods

| Software and algorithms | Source | Identifier |
| --- | --- | --- |
| MirGeneDB | Fromm et al. 2021 | doi.org/10.1093/nar/gkab1101 |
| mafft-xinsi v7.475 | Katoh et al., 2019 | doi.org/10.1093/bib/bbx108 |
| HMMER (esl-weight) | Wheeler and Eddy, 2013 | doi.org/10.1093/bioinformatics/btt403 |
| RNAalifold v2.4.17 | Lorenz et al., 2011 | doi.org/10.1186/1748-7188-6-26 |
| Infernal 1.1.4 | Nawrocki, E.P., and Eddy, S.R. 2013 | doi.org/10.1093%2Fbioinformatics%2Fbtt509 |
| cmsearch | Nawrocki, E.P., and Eddy, S.R. 2013 | doi.org/10.1093%2Fbioinformatics%2Fbtt509 |
| cmcalibrate | Nawrocki, E.P., and Eddy, S.R. 2013 | doi.org/10.1093%2Fbioinformatics%2Fbtt509 |
| Covariance models (CM) | Eddy, S.R., and Durbin, R. 1994. | doi.org/10.1093/nar/22.11.2079 |
| Snakemake v6.10.0 | Mölder et al. 2021 | f1000research.com/articles/10-33/v1 |
| MirMachine v0.2.11.2022 | This study | github.com/sinanugur/MirMachine |
| MirMachine workflow | This study | Supplementary figure 2 |
| MirMachine CM models | This study | github.com/sinanugur/MirMachine/tree/master/mirmachine/meta/cms |
| MirMachine web app | Trondsen, 2022 | https://mirmachine.org |
| MirMachine prediction<br>GFF files | This study | github.com/sinanugur/MirMachine-supplementary/tree/main/results |
| MirMachine figure data | This study | github.com/sinanugur/MirMachine-supplementary/tree/main/tables |
| R language | R Core Team<br>2022 | https://cran.r-project.org/ |
| R-chie | Lai et al. 2012 | [doi:10.1093/nar/gks241](https://doi.org/10.1093/nar/gks241) |

### Creation of high-quality CMs

MicroRNA precursor sequences were downloaded from MirGeneDB as FASTA files. We separated them into separate files based on microRNA family and we then aligned each microRNA family using the *mafft* v7.475 aligner (mafft-xinsi)^72^ and created multiple sequence alignments (MSAa) of microRNA families. We chose *mafft* since it considers secondary structure. We filtered out identical or highly similar sequences using the *esl-weight* v0.48 tool (-f --idf 0.90 --rna) from HMMER package^73^ to reduce bias due to overrepresentation of highly similar sequences. RNAalifold also expects non-identical sequences. The secondary structures of the MSAs were predicted by RNAalifold v2.4.17 (-r --noPS)^74^. Lastly, CMs for each microRNA family were generated (*cmbuild*) and calibrated (*cmcalibrate*) using Infernal^75^ and the default setting. *Cmcalibrate* is a necessary step to calibrate E-value parameters of CMs. We used the same workflow to create deuterostome and protostome specific CMs. In short, the MirGeneDB FASTA sequences were subsetted for deuterostome and protostome species.

### Determining accuracy of MirMachine predictions

First, we used the *cmsearch* function of Infernal to predict microRNA regions. In this study, true positives (TPs) are correctly predicted microRNA families and false positives (FPs) are false predictions. False negatives (FNs) refer to microRNA annotations available in MirGeneDB but not predicted by MirMachine. Using MirGeneDB and MirMachine, we extracted all true positives, false positives, and false negative predictions. We can calculate an approximation to the Matthews correlation coefficient (MCC) by using the geometric mean of sensitivity and precision. This metric is sensitive to both false negatives and false positives.

A standard *cmsearch* run reports bit score value of each prediction, which is a statistical indicator measuring the quality of an alignment score. We determined an optimal bit score value for each microRNA family to maximize MCC scores. We then filtered any MirMachine hits lower than the optimal cut-off points. We reported MCC values (and other metrics) before and after filtering. See Supplementary figure 3 for an overview of MirMachine training workflow.

**Supplementary Figure 6.**
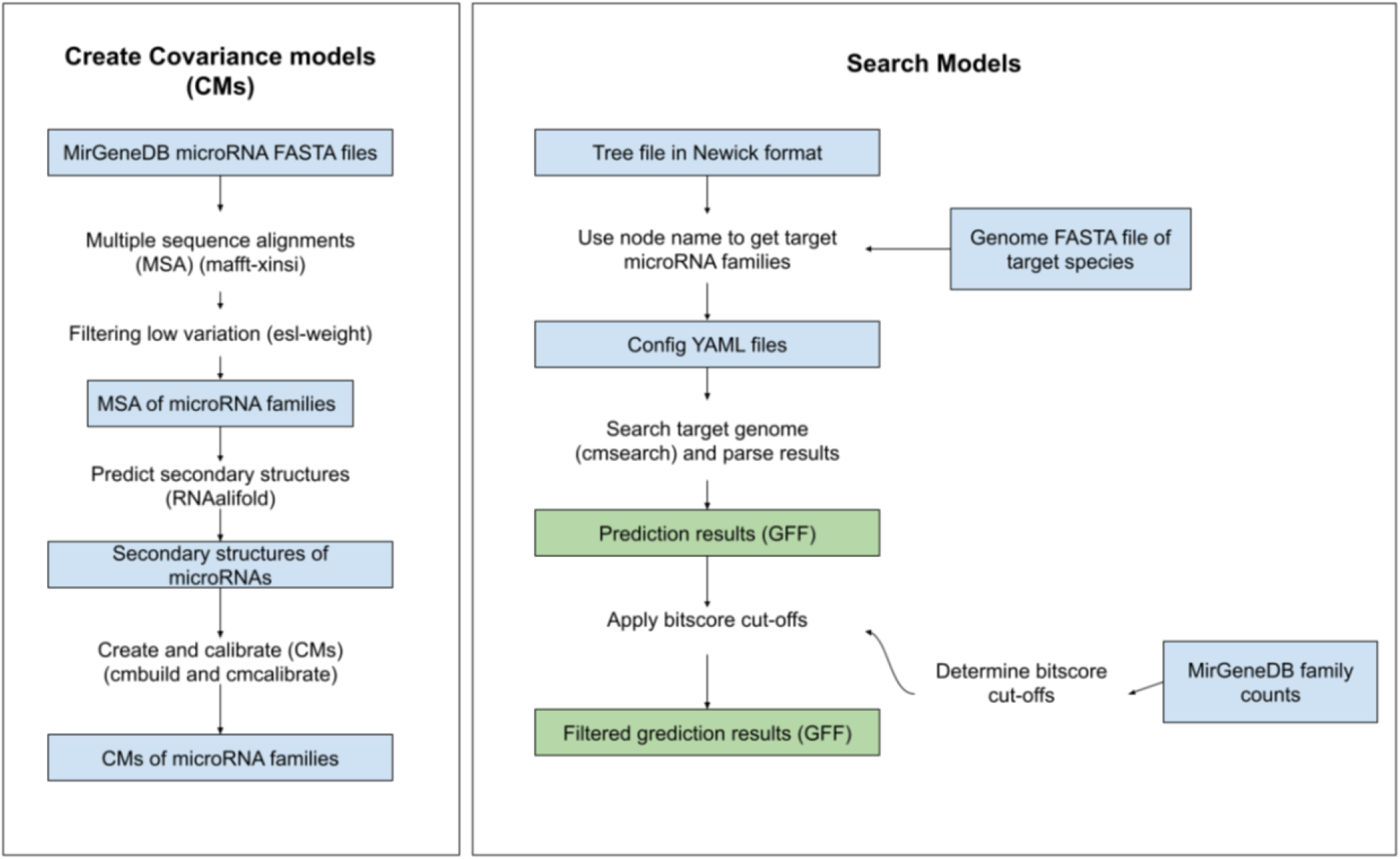
A summary of MirMachine workflow: high-quality CMs were generated using Infernal based on MirGeneDB v2.1 microRNA families. Bitscore cut-offs were determined using MirGeneDB to maximize MCC scores. We use the cutoffs to filter out low quality predictions.

### Benchmarking MirMachine models

We retrained MirMachine CM models by excluding two species: *Homo sapiens* and *Capitella teleta* and compared MirMachine performance on these species. Another benchmarking was done using Rfam models. We downloaded all microRNA models (523 in total) from the Rfam database (v 14)^32^. We predicted microRNA families using Rfam models and compared their model performance with MirMachine on selected families (e.g. LET-7, MIR-1, MIR-71, MIR-196). These families were selected because they are highly conserved and contain low false-positives or false negatives in Rfam. We also reported the total number of microRNA predictions done by both methods.

### MirMachine command line (CLI) tool

The main MirMachine engine was written in Snakemake ^76^ and the CLI wrapper in Python and R. The documentation of the MirMachine CLI tool is available at our GitHub repository. It is also available as a BioConda package ^77^ for easy installation.

### MirMachine WebApplication implementation

We implemented the web application using a software stack primarily composed of Django, React and Nginx. The application wraps the MirMachine CLI tool to provide a simpler, interactive interface for users. It is hosted at the Norwegian Research and Education Cloud (NREC), utilizing their sHPC (shared High Performance Computing) resources ^78^. It is available at https://mirmachine.org.

### Available Genome Assemblies

Lists of reference genomes of invertebrates, vertebrate mammalians and other vertebrates were downloaded from NCBI GenBank on 1/24/2022 ^79^. Analysis of yearly submitted reference genomes was conducted using Python and customized scripts.

### Covariance Model based structure plots

The Covariance Model based plots were generated using the R4RNA-package in R-chie^80^ run on R Studio version 4.2.0. The arc diagrams along with the grid-based alignment, were created with a multiple sequence alignment of all respective microRNA family members and its corresponding secondary structure as input. Within the R4RNA package, covariation was plotted, and the arc was colored based on the conservation status relative to the multiple sequence alignment provided.

### Whole genome alignment comparisons

Multiple genome alignment of 470 mammals generated with multiz as described in Hecker et al.^81^, which was kindly provided by Michael Hiller (available at http://hgdownload.soe.ucsc.edu/goldenPath/hg38/multiz470way/), was intersected with human microRNA annotations from MirGeneDB.

## Acknowledgement

B.F. is supported by the Tromsø Research foundation (Tromsø forskningsstiftelse, TFS) [20_SG_BF ‘MIRevolution’] and the UiT Aurora Outstanding program 2020-2022. S.U.U and T.B.R were supported by the Research Council of Norway under the Program Human Biobanks and Health Data (grant numbers 229621/H10 and 248791/H10). We are grateful to Michael Hiller for help with intersecting MirGeneDB with the 470 MULTIZ-alignment. We thank Wenjing Kang for help with establishing the banner plots and Eirik Høye for structure heatmap. We are grateful to Love Dalén and David Diez for help with Mammoth genomes. We would like to thank Fergal Martin and Leanne Haggerty (Ensembl), Terence Murphy (Refseq), Mark Blaxter (Darwin Tree of Life), Blake Sweeney (RNAcentral, Rfam) for discussion on the integration of MirMachine into their services and useful comments. We would like to acknowledge Torbjørn Rognes and Eivind Hovig for administrative help and we are grateful to Norwegian Research and Education Cloud (NREC) for hosting MirMachine.org.

